# Comment on “Two types of asynchronous activity in networks of excitatory and inhibitory spiking neurons”

**DOI:** 10.1101/017798

**Authors:** Rainer Engelken, Farzad Farkhooi, David Hansel, Carl van Vreeswijk, Fred Wolf

**Author notes:** Joint first authorship (alphabetically ordered) (Rainer Engelken, Farzad Farkhooi). Joint senior authorship (alphabetically ordered) (David Hansel, Carl van Vreeswijk, Fred Wolf).

## Abstract

Slow neural dynamics are believed to be important for behaviour, learning and memory [Churchland et al., 2007, Fee and Goldberg 2011]. Rate models operating in the chaotic regime show a rich dynamics at the scale of hundreds of milliseconds and provide remarkable learning capabilities [Sussillo and Abbott, 2009, Toyoizumi and Abbott, 2011, Barak et al., 2013]. However, neurons in the brain communicate via spikes and it is a major challenge in computational neuroscience to obtain similar slow rate dynamics in networks of spiking neuron models

This question was recently addressed in a paper by Ostojic (2014) published in Nature Neuroscience [Ostojic, 2014]. It argues that an “unstructured, sparsely connected network of model spiking neurons can display two fundamentally different types of asynchronous activity”. When the synaptic strength is increased, two-population (excitatory, inhibitory) networks of leaky integrate-and-fire (LIF) neurons would undergo a *phase transition* from the “well-studied asynchronous state, in which individual neurons fire irregularly at constant rates” to another asynchronous state in which “the firing rates of individual neurons fluctuate strongly in time and across neurons” [Ostojic, 2014]. These two regimes would differ in an essential manner, the rate dynamics being chaotic beyond the phase transition. As indicated by this paper’s title, the transition is its central result. Finding a transition to chaotic slow-varying rate dynamics in spiking networks in such a simple model would be an important step towards an understanding of the computations underlying behaviour and learning and fill agap in the current understanding of network dynamics. Unfortunately, we found that there is no such phase transition to chaos in the spiking network analyzed in [Ostojic, 2014]. Here, we provide simple evidence that the paper is factually incorrect and conceptually misleading.

The paper [Ostojic, 2014] starts with simulations of a network of LIF neurons for different values of the synaptic strength, J, while all other parameters are fixed to a specific set. It is observed that the population mean firing rate of the neurons, *v*_0_, is well described by a mean field calculation only below a certain coupling strength J*. At this value, the average firing rate starts to deviate from the mean field prediction more than 5%. (Fig.1a in [Ostojic, 2014], denoted Fig.P1a; hereafter figures in [Ostojic, 2014] are denoted by their numbers preceded by a “P”). In [Ostojic, 2014], it was claimed that the “classical” asynchronous state exhibit an instability at J=J*. Above J* the dynamics would still be asynchronous, but in a way which would be essentially different from the “classical” asynchronous state. To assess this claim, the author replaces the full dynamics of the spiking LIF network by a corresponding rate model, the “Poisson network”. Simulations indicate that as J increases, there is a value, J=J_c_, at which the dynamics of the latter undergo a phase transition between a state in which the rates are constant in time (fixed point) and a state in which they fluctuate chaotically with long time-scales. The author then derives an equation for J_c_ which is in agreement with the simulations of the Poisson model. For the parameters used in Fig.P1 and P2 the value of J_c_ is rather close to J*. Relying on this similarity, the paper [Ostojic, 2014] concludes that: (i) in the LIF network an instability occurs near J* which is of the same nature as the one occurring at J_c_ in the Poisson network. (ii) The asynchronous states below and above J* are essentially different in the LIF network.

**Figure 1:**
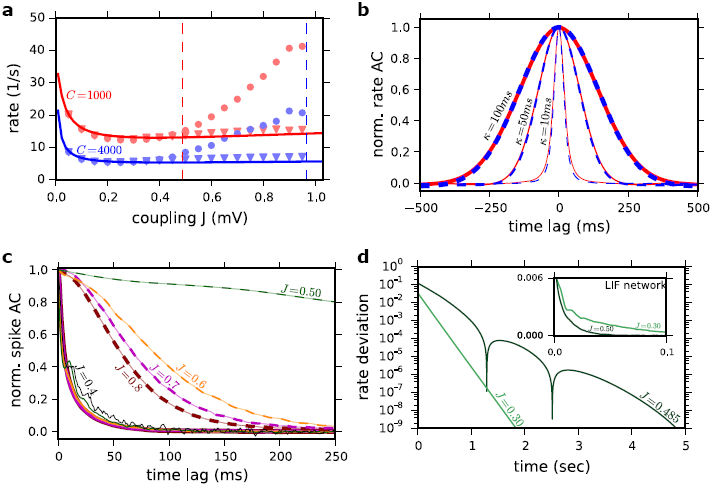
**(a)** Population averaged firing rate in the network vs. interaction strength, J. Solid lines: Ricciardi mean field for C=1000 (red) and C=4000 (blue). Predictions for J_c_ (Eq 16) are indicated by the corresponding dashed vertical lines. Simulation results (event-based simulation implemented in Julia programing language) are also plotted. Dots: Δ=0.55 ms. Triangles: Δ=0.0 ms. Results for C=1000, N=10000 (red dots) and C=4000 and N=40000 (blue dots). **(b)** Averaged normalized AC of neuronal rate functions for J=0.8 mV and C=1000 (red) and C=4000 (dashed blue) LIF networks. The rate functions were computed by filtering the spike trains of the neurons (time bin =1 ms) with a Gaussian filter with 10 ms (the thinnest lines), 50 ms (moderated lines) and 100 ms (the thickest lines) standard deviation. **(c)** Autocorrelation function of the spike trains (no filtering) normalized to the second pick. Solid lines: LIF network. Dashed lines: Poisson network. The results are shown for J = 0.5 mV (dark green), J=0.6 mV (dark orange), J = 0.7 mV (magenta) and J=0.8 mV (dark red). For the LIF the AC is also shown for J=0.4 mV (solid black). To compute the ACs for the Poisson network we simulated a network for 100 s (time step 1ms) and averaged the results over 40 realizations of the initial conditions. The network size is N=100000 for 0.5=<J=<0.6 mV and N=10000 for J>0.6 mV. For the LIF network we averaged spike autocorrelation of 3000 randomly chosen neurons with a 1 ms bin following Eq. 23 in the paper. All parameters are as in Fig.P3. **(d)** Subcritical behaviour of the systems. The absolute value of normalised response to eigenvector corresponds to λ_*max*_ in Eq. 16 in the paper for J=0.30 mV (light green lines) and J=0.485 mV (dark green lines). The main plot is reflecting the decay of this perturbation in the rate network and the inset exhibits the response (normalized to background) of spiking network (averaged over about 1.5 millions trials). The perturbation vector along the least stable direction correspond to Eq. 16 in the paper (with the norm 10) in the LIF network was applied to the constant feedforward input for 2 ms. The perturbation magnitude is adjusted to induce roughly the same response onset in rate network. The rate deviation in y-axis is |1 - 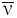| where 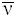 is the weighted response along the perturbation vector and it is normalised to the background.

However, as we now show, the reported agreement between the predicted transition at J_c_ and the spiking network simulation results is coincidental and due to the chosen parameters used in the paper [Ostojic, 2014] and not generally valid. We start by providing two counter-examples.

Our first counter-example is the LIF model considered in [Ostojic, 2014], we take N=40000 and C=4000 synapses per neuron instead of N=10000 and C=1000 (all other parameters as in Fig.P1, except for the network size, keeping the connection probability constant). The population firing rate, *v*_0_ (J), is plotted in Fig.1a. It deviates from the mean field prediction at J*≅0.3 mV by more than 5%. Nonetheless, the critical point in the corresponding Poisson rate network is J_c_≅ 0.96 mV and thus it is more than three times larger than J*.

Our second counter-example is the LIF network of Fig.P1 and P2 with the same parameters except for the delay, Δ: we take Δ = 0. The spiking network shows no longer a large deviation from the mean-field prediction (Fig.1a). However, the proposed analogy with the Poisson rate network still predicts that a deviation should occur at J*≅0.49 mV, since the transition to chaos in the Poisson network is independent of the delay. The author seems to be well aware of this discrepancy. Indeed, it is stated in the Online Methods that delays must be larger than the refractory period, because “if the delays are shorter, spikes that reach a neuron while it is refractory do not have an effect and the overall coupling is effectively reduced” [Ostojic, 2014]. If this were correct, this effective reduction should be reflected in the formula for predicting J_c_ (Eq. 16). This is not the case: the latter does not depend on Δ.

It is also argued in the paper [Ostojic, 2014] that the results plotted in Fig.P3a and b support the analogy between the rate dynamics of the Poisson model and the dynamics of the LIF network. However, the comparison made in this figure is conceptually misleading. In the Poisson model, the rate as a function of time is an *unequivocally* defined quantity. It is the *dynamical* variable of the model and the time scale over which the rate fluctuates for strong enough coupling is fully determined by these dynamics. This is not the case in the LIF model where the “rate” and its “dynamics” depend on the temporal width over which the spiking activity is filtered. The width of the Gaussian filter used in [Ostojic, 2014] is 50ms. This choice is arbitrary and therefore the similarity is observed in the rate autocorrelations (ACs) plotted in the upper and lower panels in Fig.P3b which depends on this choice (Fig.1b). Moreover, the spike ACs plotted in Fig. P3c for the two models exhibit essential differences as we now show.

For J=0.2 and 0.4 mV, the spike AC in the Poisson rate model (Fig.P3c, upper panel) is close to a Dirac function reflecting that the dynamics is at fixed point - that is the rate variable from which the Poisson process of the spikes is generated is constant. For J=0.6 mV the spike AC is very different: a broad component has now appeared. It is flat at zero time lag and has a negative curvature at short time lags (Fig.1c and Fig.P3c). A detailed analysis reveals that this change has all the characteristics of a true phase transition. It shows that close to the phase transition, the amplitude vanishes proportionally to J-J_c_ and the decorrelation time diverges as 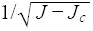 (Fig.S1a).

The spike AC behaves very differently in the LIF network. For J=0.2 mV it exhibits at zero time lag a sharp peak flanked by a trough which reflects the refractoriness (absolute and relative) of the single neuron dynamics. As J increases, there is a progressive changes in the AC shape. Eventually, the trough disappears. The flanks of the zero peak are now decreasing exponentially (Fig.1c, solid lines). A careful analysis shows that the typical time constant of this decrease depends only weakly on J (Fig.1c, solid lines). It is always on the order of the membrane time constant of the neurons (20ms). Note also that by contrast with what is observed in the Poisson network, for J=0.5 to 0.8 mV, the spike AC curvature is always positive and peaked around zero time lag (Fig.1c, dashed lines).

Additionally, in order to compare the behaviour of both models below the postulated phase transition, we perturbed rate and spiking networks of identical topology in the least stable direction, predicted by Eq. 16 in the paper [Ostojic, 2014]. Fig.1d shows that the decay of the perturbation in the rate network slows down near the transition, indicating a critical slowing down (Fig.1d, main panel). If there were an analogous transition in the spiking network, also its perturbation should decay slower as the transition is approached. Our result (Fig.1d, inset) shows that the decay time-scales of the perturbation is independent of J and it stays close to the membrane time constant (similar to solid lines in Fig.1c).

We therefore conclude that, contrary to what is argued, the spiking LIF network studied in [Ostojic, 2014] does not exhibit a phase transition to a chaotic state similar to the one occurring in the studied rate model. The reported mismatch between the average firing rate in this LIF network simulations and the mean-field calculation is unrelated to such a transition. Understanding the conditions under which slow rate dynamics are generated in *spiking neural networks* remain an open challenge.

## Acknowledgments

We thank Ran Darshan, Omri Harish and Gianluigi Mongillo for fruitful discussions. The work of DH and CvV was partially supported by grants ANR-13-BSV4-0014-03 - BALAV1 and ANR-14-NEUC-0001-01-BASCO and performed in the framework of the France-Israel Laboratory of Neuroscience (FILN). FF was supported by the BMBF, FKZ 01GQ 1001B and GIF-I-1224-396.13/2012. RE and FW received funding from Evangelisches Studienwerk Villigst, DFG through CRC 889 and Volkswagen Foundation.

## Supplement figure caption

**Figure S1:**
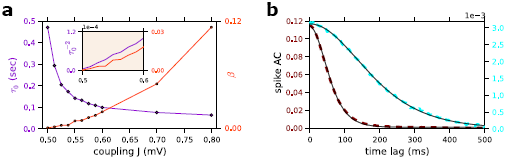
**(a)** The decorrelation time (τ_0_, violet diamond, left y-axis) and amplitude at zero time lag (beta, orange circles, right y-axis) of the baseline-subtracted population averaged spike AC are plotted vs. J for the Poisson network. These parameters were obtained by fitting the spike AC with *ACF* (τ) = β/*cosh*(τ/τ_0_)^2^ (see Fig.S1b). Inset: the rescaled estimated -2 (left axis, violet) and β values (orange, right axis) for J=0.5, 0.5125, 0.525, 0.5375, 0.55, 0.5625,0.575, 0.5875 and 0.6 mV, to show that they vanish linearly near the phase transition. **(b)**The non-normalized spike AC can be very well fitted with *ACF* (τ) = β/*cosh*(τ/τ_0_)^2^. Dashed lines: Simulation results; J=0.525 mV (cyan, right y-axis) and J=0.8 mV (dark red, left y-axis): Black solid line: The fits. The parameters of the fit are used in Fig1.D in the main text.

